# ASA^3^P: An automatic and scalable pipeline for the assembly, annotation and higher level analysis of closely related bacterial isolates

**DOI:** 10.1101/654319

**Authors:** Oliver Schwengers, Andreas Hoek, Moritz Fritzenwanker, Linda Falgenhauer, Torsten Hain, Trinad Chakraborty, Alexander Goesmann

## Abstract

Whole genome sequencing of bacteria has become daily routine in many fields. Advances in DNA sequencing technologies and continuously dropping costs have resulted in a tremendous increase in the amounts of available sequence data. However, comprehensive in-depth analysis of the resulting data remains an arduous and time consuming task. In order to keep pace with these promising but challenging developments and to transform raw data into valuable information, standardized analyses and scalable software tools are needed. Here, we introduce ASA^3^P, a fully automatic, locally executable and scalable assembly, annotation and analysis pipeline for bacterial genomes. The pipeline automatically executes necessary data processing steps, *i.e.* quality clipping and assembly of raw sequencing reads, scaffolding of contigs and annotation of the resulting genome sequences. Furthermore, ASA^3^P conducts comprehensive genome characterizations and analyses, *e.g.* taxonomic classification, detection of antibiotic resistance genes and identification of virulence factors. All results are presented via an HTML5 user interface providing aggregated information, interactive visualizations and access to intermediate results in standard bioinformatics file formats. We distribute ASA^3^P in two versions: a locally executable Docker container for small-to-medium-scale projects and an OpenStack based cloud computing version able to automatically create and manage self-scaling compute clusters. Thus, automatic and standardized analysis of hundreds of bacterial genomes becomes feasible within hours. The software and further information is available at: http://asap.computational.bio.

## Introduction

In 1977 DNA sequencing was introduced to the scientific community by Frederick Sanger [1]. Since then, DNA sequencing has come a long way from dideoxy chain termination over high-throughput sequencing of millions of short DNA fragments and finally to real-time sequencing of single DNA molecules [2,3]. Latter technologies of so called next generation sequencing (NGS) and third generation sequencing have caused a massive reduction of time and costs, and thus, led to an explosion of publicly available genomes. In 1995, the first bacterial genomes of *M. genitalium* and *H. influenzae* were published [4,5]. Today, the NCBI RefSeq database release 93 alone contains 54,854 genomes of distinct bacterial organisms [6]. Due to the maturation of NGS technologies, the laborious task of bacterial whole genome sequencing (WGS) has transformed into plain routine [7] and nowadays, has become feasible within hours [8].

As the sequencing process is not a limiting factor anymore, focus has shifted towards deeper analyses of single genomes and also large cohorts of *e.g.* clinical isolates in a comparative way to unravel the plethora of genetic mechanisms driving diversity and genetic landscape of bacterial populations [9]. Comprehensively characterizing bacterial organisms has become a desirable and necessary task in many fields of application including environmental- and medical microbiology [10]. The recent worldwide surge of multi-resistant microorganisms has led to the realization, that without the implementation of adequate measures in 2050 up to 10 million people could die each year due to infections with antimicrobial resistant bacteria alone [11]. Thus, sequencing and timely characterization of large numbers of bacterial genomes is a key element for successful outbreak detection, proper surveillance of emerging pathogens and monitoring the spread of antibiotic resistance genes [12]. Comparative analysis could lead to the identification of novel therapeutic drug targets to prevent the spread of pathogenic and antibiotic-resistant bacteria [13–16].

Another very promising and important field of application for microbial genome sequencing is modern biotechnology. Due to deeper knowledge of the underlying genomic mechanisms, genetic engineering of genes and entire bacterial genomes has become an indispensable tool to transform them into living chemical factories with vast applications, as for instance, production of complex chemicals [17], synthesis of valuable drugs [18–20] and biofuels [21], decontamination and degradation of toxins and wastes [22,23] as well as corrosion protection [24].

Now, that the technological barriers of WGS have fallen, genomics finally transformed into Big Data science [25] inducing new issues and challenges [26]. To keep pace with these developments, we believe that continued efforts are required in terms of the following issues:

a. Automation: Repeated manual analyses are time consuming and error prone. Following the well known “don’t repeat yourself” mantra and the pareto principle, scientists should be able to concentrate on interesting and promising aspects of data analysis instead of ever repeating data processing tasks.
b. Standard operating procedures (SOPs): In a world of high-throughput data creation and complex combinations of bioinformatic tools, SOPs are indispensable to increase and maintain both reproducibility and comparability [27].
c. Scalability: To keep pace with available data, bioinformatics software needs to take advantage of modern compute technologies, *e.g.* multi-threading and cloud computing.

Addressing these issues, several major platforms for the automatic annotation and analysis of prokaryotic genomes have evolved in recent years as for example the NCBI Prokaryotic Genome Annotation Pipeline [6], RAST [28] and PATRIC [29]. All three provide sophisticated genome analysis and annotation pipelines and pose a de-facto community standard in terms of annotation quality. In addition, several offline tools, *e.g.* Prokka [30], have been published in order to address major drawbacks of the aforementioned online tools, *i.e.* they are not executable on local computers or in on-premises cloud computing environments. However, comprehensive analysis of bacterial WGS data is not limited to the process of annotation alone but also requires sequencing technology-dependent pre-processing of raw data as well as subsequent characterization steps. As analysis of bacterial isolates and cohorts will be a standard method in many fields of application in the near future, demand for sophisticated local assembly, annotation and higher-level analysis pipelines will rise constantly. Furthermore, we believe that the utilization of portable devices for DNA sequencing will shift analysis from central software installations to either decentral offline tools or scalable cloud solutions. To the authors’ best knowledge, there is currently no published bioinformatics software tool successfully addressing all aforementioned issues. In order to overcome this bottleneck, we introduce ASA^3^P, an automatic and scalable software pipeline for the assembly, annotation and higher level analysis of closely related bacterial isolates.

## Design and implementation

ASA^3^P is implemented as a modular command line tool in Groovy (http://groovy-lang.org), a dynamic scripting language for the Java virtual machine. In order to achieve acceptable to best possible results over a broad range of bacterial genera, sequencing technologies and sequencing depths, ASA^3^P incorporates and takes advantage of published and well performing bioinformatics tools wherever available and applicable in terms of lean and scalable implementation. As the pipeline is also intended to be used as a preprocessing tool for more specialized analyses, it provides no user-adjustable parameters by design and thus facilitates the implementation of robust SOPs. Hence, each utilized tool is parameterized according to community best practices and knowledge (S1 Table).

### Workflow, tools and databases

Depending on the sequencing technology used to generate the data, ASA^3^P automatically chooses appropriate tools and parameters. Semantically, the pipeline’s workflow is divided into four stages (Fig 1). In the first stage A (Fig 1A), provided input data are processed, resulting in annotated genomes. Therefore, raw sequencing reads are quality controlled and clipped via FastQC (https://github.com/s-andrews/FastQC), FastQ Screen (https://www.bioinformatics.babraham.ac.uk/projects/fastq_screen), Trimmomatic [31] and Filtlong (https://github.com/rrwick/Filtlong). Filtered reads are then assembled via SPAdes [32] for Illumina reads, HGAP 4 [33] for Pacific Bioscience (PacBio) reads and Unicycler [34] for Oxford Nanopore Technology (ONT) reads, respectively. Hybrid assemblies of Illumina and ONT reads are conducted via Unicycler, as well. Before annotating assembled genomes with Prokka [30], contigs are rearranged and ordered via the multi-reference scaffolder MeDuSa [35]. For the annotation of subsequent pseudogenomes ASA^3^P uses custom genus-specific databases based on binned RefSeq genomes [6] as well as specialized protein databases, *i.e.* CARD [36] and VFDB [37]. In order to integrate public or externally analyzed genomes, ASA^3^P is able to incorporate different types of pre-processed data, *e.g.* contigs, scaffolds and annotated genomes.

**Fig 1.**
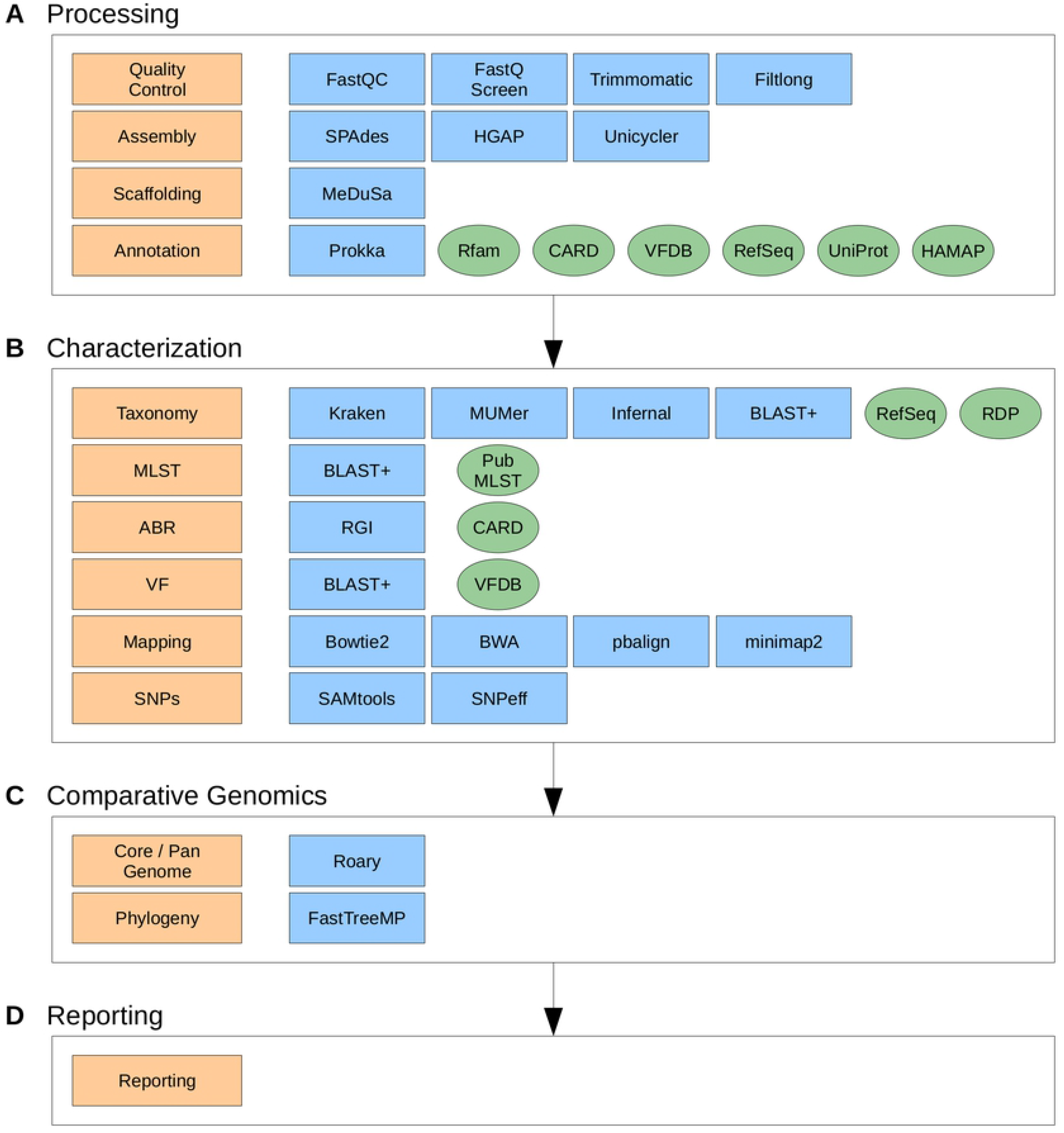
Workflow stages, analysis steps and incorporated third party executables and databases. Internal analysis steps (orange boxes) are organized in four stages (large white boxes, A-D). Each step takes advantage of third party executables (blue boxes) and/or databases (green ovals) depending on input data types.

In a second stage B (Fig 1B), all assembled and annotated genomes are extensively characterized. A taxonomic classification is conducted comprising three distinct methods, *i.e.* a kmer profile search, a 16S sequence homology search and a computation of an average nucleotide identity (ANI) [38] against user provided reference genomes. For the kmer profile search, the software takes advantage of the Kraken package [39] and a custom reference genome database based on RefSeq [6]. The 16S based classification is implemented using BLAST+ [40] and the RDP [41] database. Calculation of ANI values is implemented in Groovy using nucmer within the MUMmer package [42]. A subspecies level multi locus sequence typing (MLST) analysis is implemented in Groovy using BLAST+ [40] and the PubMLST.org [43] database. Detection of antibiotic resistances (ABRs) is conducted via RGI and the CARD [36] database. A detection of virulence factors is implemented via BLAST+ [40] and VFDB [37]. Quality clipped reads get mapped onto user provided reference genomes via Bowtie2 [44] for Illumina, pbalign (https://github.com/PacificBiosciences/pbalign) for PacBio and Minimap2 [45] for ONT sequence reads, respectively. Based on these read mappings, the pipeline calls, filters and annotates single nucleotide polymorphisms (SNPs) via SAMtools [46] and SnpEff [47] and finally computes consensus sequences for each isolate. In order to maximize parallel execution and thus reducing overall runtime, stage A and B are technically implemented as a single step.

A third comparative stage C (Fig 1C) is triggered as soon as stages A and B are completed, *i.e.* all genomes are processed and characterized. Utilizing aforementioned consensus sequences, ASA^3^P computes a phylogenetic approximately maximum-likelihood tree via FastTreeMP [48]. This is complemented by the calculation of a core, accessory and pan-genome as well as the detection of isolate genes conducted via Roary [49].

In a final stage (Fig 1D), the pipeline aggregates all analysis results and data files and finally provides a graphical user interface (GUI), *i.e.* responsive HTML5 documents comprising detailed information via interactive widgets and visualizations. Therefore, ASA^3^P takes advantage of modern web frameworks, *e.g.* Bootstrap (https://getbootstrap.com) and jQuery (https://jquery.com) as well as adequate JavaScript visualization libraries, *e.g.* Google Charts (https://developers.google.com/chart), D3 (https://d3js.org) and C3 (http://c3js.org).

### User input and output

Each set of bacterial isolates to be analyzed within a single execution is considered as a self-contained analysis of bacterial cohorts and is subsequently referred to as an ASA^3^P project. As ASA^3^P was developed in order to analyze cohorts of closely related isolates, *e.g.* a clonal outbreak, the pipeline expects all genomes within a project to belong to at least the same genus, although a common species is most favourable. For each project, the pipeline expects a distinct directory comprising a configuration spreadsheet containing necessary project information and a subdirectory containing all input data files. Such a directory is subsequently referred to as project directory. In order to ease provisioning of necessary information, we provide a configuration spreadsheet template comprising two sheets (S1 and S2 Figs). The first sheet contains project meta information such as project names and descriptions as well as contact information on project maintainers and provided reference genomes. The second sheet stores information on each isolate comprising an unique identifier as well as data input type and related files. ASA^3^P is currently able to process input data in the following standard file formats: Illumina paired-end and single-end reads as compressed FastQ files, PacBio RSII and Sequel reads provided either as single unmapped bam files or via triples of bax.h5 files, demultiplexed ONT reads as compressed FastQ files, pre-assembled contigs or pseudogenomes as Fasta files and pre-annotated genomes as Genbank, EMBL or GFF files. In the latter case, corresponding genome sequences can either be included in the GFF file or provided via separate Fasta files.

As ASA^3^P is also intended to be used as an automatic preprocessing tool providing as much reliable information as possible, results are stored in a standardized manner within project directories comprising quality clipped reads, assemblies, ordered and scaffolded contigs, annotated genomes, mapped reads, detected SNPs as well as ABRs and virulence factors. In detail, all result files are stored in distinct subdirectories for each analysis by the pipeline and for certain analyses further subdirectories are created therein for each genome (S3 Fig). Aggregated information is stored in a standardized but flexible document structure as JSON files. Text and binary result files are stored in standard bioinformatics file formats, *i.e.* FastQ, Fasta, BAM, VCF and Newick. Providing results in such a machine-readable manner, ASA^3^P outputs can be further exploited by manual or automatic downstream analyses since customized scripts with a more targeted focus can easily access necessary data. In addition, ASA^3^P creates user-friendly HTML5 reports providing both prepared summaries as well as detailed information via sophisticated interactive visualizations.

### Implementation and software distributions

ASA^3^P is designed as a modular and expandable application with high scalability in mind. It consists of three distinct tiers, *i.e.* a command line interface, an application programming interface (API) and analysis specific cluster distributable worker scripts. A common software-wide API is implemented in Java whereas the core application and worker scripts are implemented in Groovy. In order to overcome common error scenarios on distributed high-performance compute (HPC) clusters and cloud infrastructures and thereby delivering robust runtime behavior, the pipeline takes advantage of a well-designed shared file system oriented data organization, following a convention over configuration approach. Thus, loosely coupled software parts run both concurrently and independently without interfering with each other. In addition, future enhancements and externally customized scripts reliably find intermediate files at reproducible locations within the file system.

As ASA^3^P requires many third party dependencies such as software libraries, bioinformatics tools and databases, both distribution and installation is a non-trivial task. In order to reduce the technical complexity as much as possible and to overcome this bottleneck for non-computer-experts, we provide two distinct distributions addressing different use cases and project sizes, *i.e.* a locally executable containerized version based on Docker (DV) (https://www.docker.com) as well as an OpenStack (OS) (https://www.openstack.org) based cloud computing version (OSCV). Details and appropriate use cases of both are described in the following sections.

#### Docker

For small to medium projects and utmost simplicity we provide a Docker container image encapsulating all technical dependencies such as software libraries and system-wide executables. As the DV offers only vertical scalability, it addresses small projects of less than ca. 200 genomes. The necessary container image is publicly available from our Docker repository (https://hub.docker.com/r/oschwengers/asap) and can be started without any prior installation, except of the Docker software itself. For the sake of lightweight container images and to comply with Docker best practices, all required bioinformatics tools and databases are provided via an additional tarball, subsequently referred to as ASA^3^P volume which users merely need to download and extract, once. For non-Docker savvy users, a shell script hiding all Docker related aspects is also provided. By this, executing the entire pipeline comes down to a single command:

~~~
*sudo <asap_volume>/asap-docker.sh <project_path>*.
~~~

#### Cloud Computing

For medium to very large projects, we provide an OS based version in order to utilize horizontal scaling capabilities of modern cloud computing infrastructures. Since creation and configuration of such complex setups require advanced technical knowledge, we provide a shell script taking care of all cloud specific aspects and to orchestrate and execute the underlying workflow logic. Necessary cloud specific properties such as available hardware quotas, virtual machine (VM) flavours and OS identifiers are specified and stored in a custom property file, once. In order to address contemporary demands for high scalability, the OSCV is able to horizontally scale out and distribute workloads on an internally managed Sun Grid Engine (SGE) based compute cluster. A therefore indispensable shared file system is provided by an internal network file system (NFS) server sharing distinct storage volumes for both project data and a necessary ASA^3^P volume. In order to create and orchestrate both software and hardware infrastructures in a fully automatic manner, the pipeline takes advantage of the BiBiGrid (https://github.com/BiBiServ/bibigrid) framework. Hereby, ASA^3^P is able to adjust the compute cluster size fitting the number of isolates within a project as well as available hardware quotas. Except of an initial VM acting as a gateway into an OS cloud project, the entire compute cluster infrastructure is automatically created, setup, managed and finally shut down by the software. Thus, ASA^3^P can exploit vast hardware capacities and is portable to any OS compatible cloud. For further guidance, all prerequisite installation steps are covered in a detailed user manual.

## Results

### Analysis features

ASA^3^P conducts a comprehensive set of pre-processing tasks and genome analyses. In order to delineate currently implemented analysis features, we created and analyzed a benchmark data set comprising 32 Illumina sequenced *Listeria monocytogenes* isolates randomly selected from SRA as well as four *Listeria monocytogenes* reference genomes from Genbank (S2 Table). All isolates were successfully assembled, annotated, deeply characterized and finally included in comparative analyses. Table 1 provides genome wise minimum and maximum values for key metrics covering results from workflow stages A and B. After conducting a quality control and adapter removal for all raw sequencing reads, a minimum of 393,300 and a maximum of 6,315,924 reads remained, respectively. Genome wise minimum and maximum mean phred scores were 34.7 and 37.2. Assembled genome sizes ranged between 2,818 kbp and 3,201 kbp with a minimum of 12 and a maximum of 108 contigs. Hereby, a maximum N50 of 1,568 kbp was achieved. After rearranging and ordering contigs to aforementioned reference genomes, assemblies were reduced to 1 to 10 scaffolds and 0 to 42 contigs per genome, thus increasing the minimum and maximum N50 to 658 kbp and 3,034 kbp, respectively. Pseudolinked genomes were subsequently annotated resulting in between 2,735 and 3,200 coding genes and between 95 and 144 non-coding genes.

**Table 1.** Per-isolate key metrics of an ASA^3^P analysis. Selected per-isolate key metrics automatically calculated by ASA^3^P analyzing a benchmark data set comprising 32 *Listeria monocytogenes* isolates.

| Analysis | Metric | Minimum | Maximum |
| --- | --- | --- | --- |
| QC | # reads | 393,300 | 6,315,924 |
| QC | Mean read length | 125.7 nt | 228.5 nt |
| QC | mean Phred score | 34.7 | 37.2 |
| assembly | Genome size | 2,817,892 bp | 3,201,054 bp |
| assembly | # contigs | 12 | 108 |
| assembly | N50 | 56,125 bp | 1,568,056 bp |
| assembly | GC content | 37% | 38% |
| scaffolding | # scaffolds | 1 | 10 |
| scaffolding | # contigs | 0 | 42 |
| scaffolding | N50 | 657,549 bp | 3,034,489 bp |
| annotation | # coding genes | 2,735 | 3,200 |
| annotation | # non-coding genes | 95 | 144 |
| antibiotic resistance | # ABR genes | 0 | 2 |
| virulence factors | # vf genes | 16 | 35 |

After pre-processing, assembling and annotating all isolates, ASA^3^P successfully conducted deep characterizations of all isolates, which were consistently classified to the species level via kmer-lookups as well as 16S ribosomal RNA database searches as *Listeria monocytogenes*, except of a single isolate classified as *Listeria innocua*. In line with these results all isolates shared an ANI value above 95% and a conserved DNA of at least 80% with at least one of the reference genomes, except for the *L. innocua* isolate which shared a maximum ANI of 90.7% and a conserved DNA of only 37.3%. Furthermore, the pipeline successfully subtyped all but one of the isolates via MLST, by automatically detecting and applying the “lmonocytogenes” schema. Noteworthy, the *L. innocua* isolate constitutes a distinct MLST lineage, *i.e. L. innocua*. ASA^3^P detected between 0 and 2 antibiotic resistance genes and between 16 and 35 virulence factor genes. A comprehensive list of all key metrics for each genome is provided in a separate spreadsheet (S1 File).

Finally, core and pan-genomes were computed resulting in 1,485 core genes and a pan-genome comprising 7,242 genes. Excluding the *L. innocua* strain and re-analyzing the dataset reduced the pan-genome to 6,197 genes and increased the amount of core genes to 2,004 additionally endorsing its taxonomic difference.

### Data visualization

Analysis results as well as aggregated information get collected, transformed and finally presented by the pipeline via user friendly and detailed reports. These comprise local and responsive HTML5 documents containing interactive JavaScript visualizations facilitating the easy comprehension of the results. Fig 2 shows an exemplary collection of embedded data visualizations. Where appropriate, specialized widgets were implemented, as for instance circular genome annotation plots presenting genome features, GC content and GC skew on separate tracks (Fig 2A). These plots can be zoomed, panned and downloaded in SVG format for subsequent re-utilization. Another example is the interactive and dynamic visualization of SNP based phylogenetic trees (Fig 2E) via the Phylocanvas library (http://phylocanvas.org) enabling customizations by the user, as for instance changing tree types as well as collapsing and rotating subtrees. In order to provide users with an expeditious but conclusive overview on bacterial cohorts, key genome characteristics are visualized via an interactive parallel coordinates plot (Fig 2F) allowing for the combined selection of value ranges in different dimensions. Thus, clusters of isolates sharing high-level genome characteristics can be explored and identified straightforward. In order to rapidly compare different ABR capabilities of individual isolates, a specialized widget was designed and implemented (Fig 2D). For each isolate an ABR profile based on detected ABR genes grouped to 34 distinct target drug classes is computed, visualized and stacked for the easy perception of dissimilarities between genomes. Throughout the reports wherever appropriate, numeric results are interactively visualized as, for instance, the distribution of detected MLST sequence types (Fig 2B) and per-isolate analysis results summarized via key metrics presented within sortable and filterable data tables (Fig 2C).

**Fig 2.**
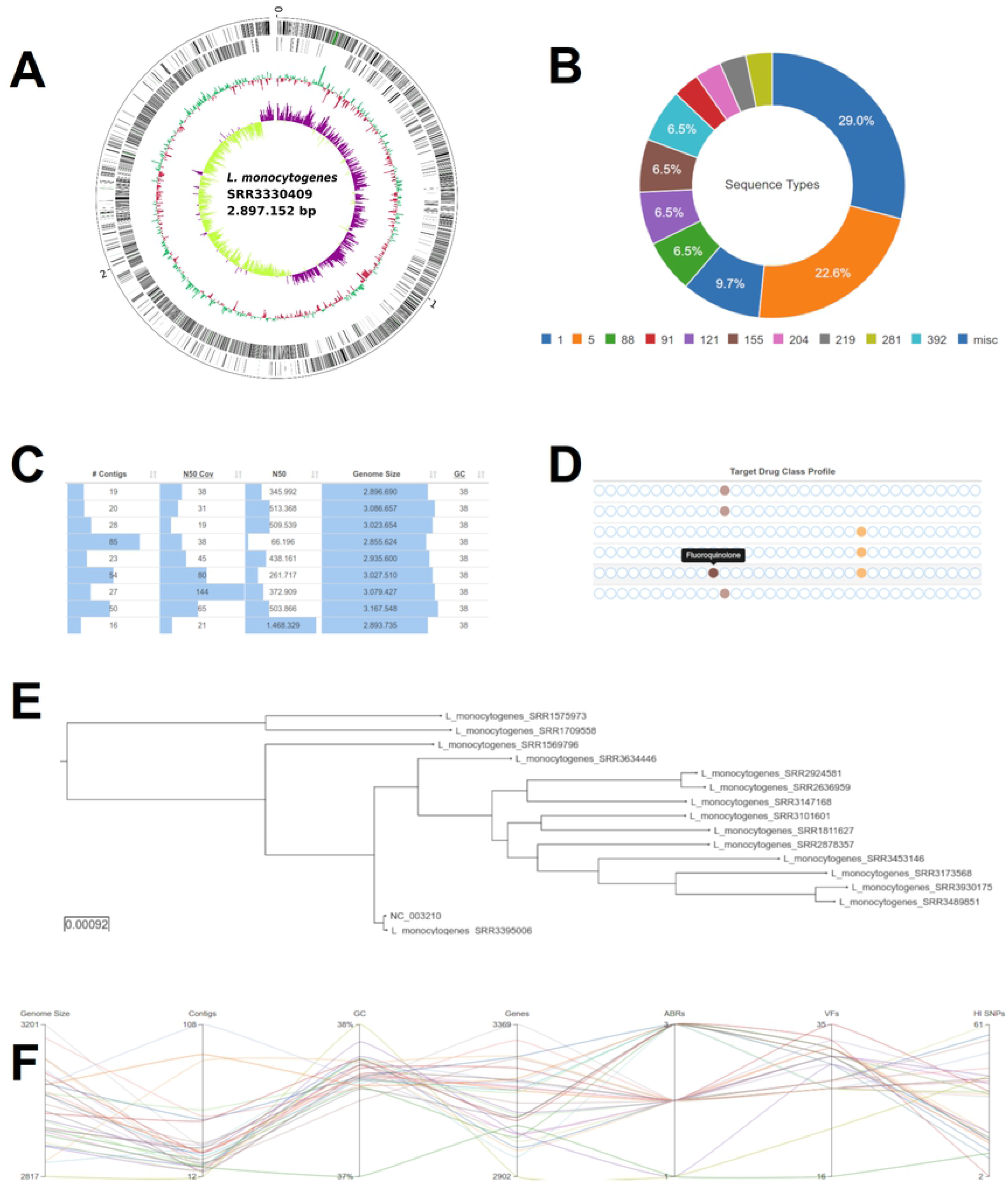
Selected screenshots of interactive GUI widgets embedded in generated HTML5 reports. (A) Circular genome plot for *Listeria monocytogenes* strain SRR3330409. From the outermost inward: genes on the forward and reverse strand, respectively, GC content and GC skew. (B) Donut chart of MLST sequence type distribution. (C) Visual representation of assembly key statistics normalized to minimum and maximum values column-wise. (D) Antibiotic resistance profiles of 6 isolates for 34 distinct target drug classes and an on-hover tooltip. (E) SNP-based phylogenetic tree. (F) Parallel coordinates plot visualizing multi-dimensional relations of key genome characteristics.

### Scalability and hardware requirements

When analyzing projects with growing numbers of isolates, local execution can quickly become infeasible. In order to address varying amounts of data, we provide two distinct ASA^3^P distributions based on Docker and cloud computing environments. Each features individual scalability properties and implies different levels of technical complexity in terms of distribution and installation requirements. In order to benchmark the pipeline’s scalability, we measured wall clock runtimes analyzing two projects comprising 32 and 1,024 *L. monocytogenes* isolates, respectively (S2 Table). Accession numbers for the large data set will be provided upon request. In addition to both public distributions, we also tested a custom installation on an inhouse SGE-based HPC cluster. The DV was executed on a VM providing 32 vCPUs and 64 GB memory. The quotas of the OS cloud project allowed for a total amount of 560 vCPUs and 1,280 GB memory. The HPC cluster comprised 20 machines with 40 cores and 250 GB memory, each. All machines hosted an Ubuntu 16.04 operating system. Table 2 shows the best-of-three runtimes for each version and benchmark data set combination. The pipeline successfully finished all benchmark analyses, except of the 1,024 dataset analyzed by the DV, due to lacking memory capacities required for the calculation of a phylogenetic tree comprising this large amount of genomes. Analyzing the 32 *L. monocytogenes* data set on larger compute infrastructures, *i.e.* the OS cloud (5:02:24 h) and HPC cluster (4:49:24 h), shows significantly reduced runtimes by approximately 50%, compared to the Docker-based executions (10:59:34 h). Not surprisingly, runtimes of the OSCV are slightly longer than HPC runtimes, due to the inherent overhead of automatic infrastructure setup and management procedures. Excluding these overheads reduces runtimes by approximately half an hour, leading to slightly shorter periods compared to the HPC version. We attribute this to a saturated workload distribution combined with faster CPUs in the cloud as stated in Table S3. Comparing measured runtimes for both data sets exhibit a ~5.8- and ~6.9-fold increase for the HPC cluster (27:56:37 h) and OSCV (34:47:45 h) version, respectively, although the amount of isolates was increased 32-fold.

**Table 2.**
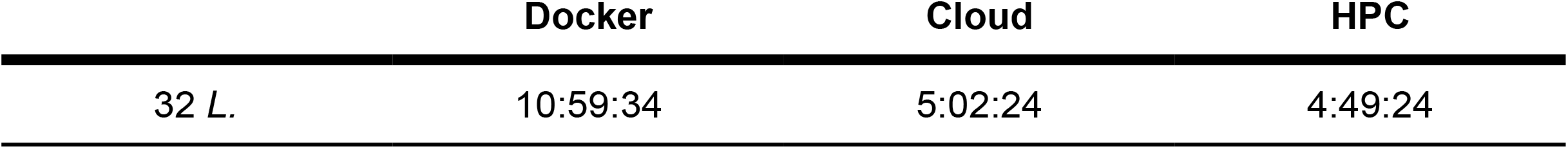
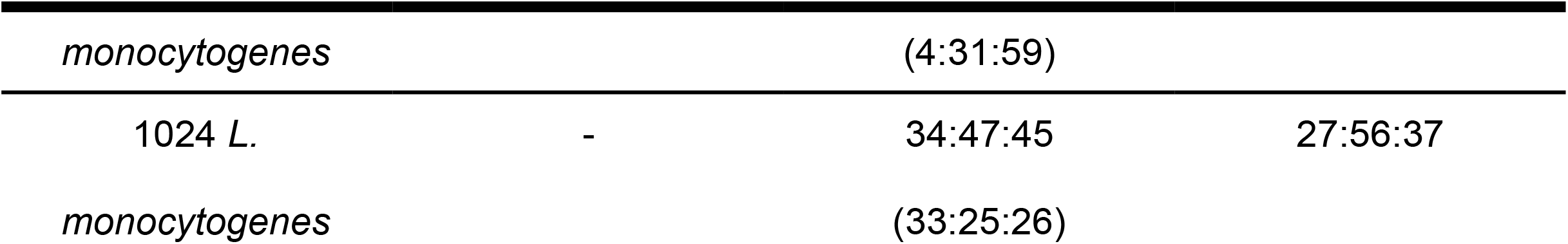
Best-of-three wall clock runtimes of ASA^3^P versions utilizing different hardware infrastructures. Wall clock runtimes times given in *hh:mm:ss* format. Docker: single virtual machine with 32 vCPUs and 64 GB memory; HPC: SGE-based HPC cluster comprising 20 nodes with 40 cores and 250 GB memory each; Cloud: OS based cloud computing project comprising 560 vCPUs and 1,280 GB memory in total, runtimes in parenthesis exclude build time for automatic infrastructure setups, *i.e.* pure ASA^3^P runtime.

|  | Docker | Cloud | HPC |
| --- | --- | --- | --- |
| 32 *L.* | 10:59:34 | 5:02:24 | 4:49:24 |

| *monocytogenes* |  | (4:31:59) |  |
| --- | --- | --- | --- |
| 1024 *L.* | - | 34:47:45 | 27:56:37 |
| *monocytogenes* |  | (33:25:26) |  |

We furthermore investigated internal pipeline scaling properties for combinations of fixed and varying HPC cluster and project sizes (S4 Fig). In a first setup, growing numbers of *L. monocytogenes* isolates were analyzed utilizing a fixed-size HPC cluster of 4 compute nodes providing 32 vCPUs and 64 GB RAM each. Iteratively doubling the amount of isolates from 32 to 1,024 led to runtimes approximately increasing by a factor of 2, in line with our expectations. Nevertheless, we observed an overproportional increase in runtime of the internal comparative steps within stage C compared to the per-isolate steps of stage A and B. We attribute this to the implementations and inherent algorithms of internally used third party executables. As this might become a bottleneck for the analysis of even larger projects, this will be subject to future developments.

In addition, we repetitively analyzed a fixed number of 128 *L. monocytogenes* isolates while increasing underlying hardware capacities, *i.e.* available HPC compute nodes. In this second setup, we could measure significant runtime reductions for up to 8 compute nodes. Further hardware capacity expansions led to saturated workload distributions and contributed negligible runtime benefits. To summarize all conducted runtime benchmarks, we conclude, that ASA^3^P is able to horizontally scale-out to larger infrastructures and thus, conducting expeditious analysis of large projects within favourable periods of time.

To test the reliable distribution and robustness of the pipeline, we executed the DV on an Apple iMac running MacOS 10.14.2 providing 4 cores and 8 GB of memory. ASA^3^P successfully analyzed a downsampled dataset comprising 4 *L. monocytogenes* isolates within a measured wall clock runtime of 8:43:12 hours. In order to assess minimal hardware requirements, the downsampled data set was analyzed iteratively reducing provided memory capacities of an OS VM. Hereby, we could determine a minimal memory requirement of 8 GB and thus draw the conclusion that ASA^3^P allows the execution of a sophisticated workflow for the analysis of bacterial WGS data cohorts on ordinary consumer hardware. However, since larger amounts of isolates, more complex genomes or deeper sequencing coverages might result in higher hardware requirements, we nevertheless recommend at least 16 GB of memory.

## Conclusion

We described ASA^3^P, a new software tool for the local, automatic and highly scalable analysis of bacterial WGS data. The pipeline integrates many common analyses in a standardized and community best practices manner and is available for download either as a local command line tool encapsulated and distributed via Docker or a self-orchestrating OS cloud version. To the authors’ best knowledge it is currently the only publicly available tool for the automatic high-throughput analysis of bacterial cohorts WGS data supporting all major contemporary sequencing platforms, offering SOPs, robust scalability as well as a user friendly and interactive graphical user interface whilst still being locally executable and thus offering on-premises analysis for sensitive or even confidential data. So far, ASA^3^P has been used to analyze thousands of bacterial isolates covering a broad range of different taxons.

## Availability and future directions

The source code is available on GitHub under GPL3 license at https://github.com/oschwengers/asap. The Docker container image is accessible at Docker Hub: https://hub.docker.com/r/oschwengers/asap. The ASA^3^P software volume containing third-party executables and databases, OpenStack cloud scripts, a comprehensive manual, configuration templates and exemplary data projects can be downloaded at http://www.computational.bio/software/asap. Questions and issues can be sent to, bug reports can be filed as GitHub issues.

Albeit ASA^3^P itself is published and distributed under a GPL3 license, some of its dependencies bundled within the ASA^3^P volume are published under different license models, *e.g.* CARD and PubMLST. Comprehensive license information on each dependency and database is provided as a DEPENDENCY_LICENSE file within the ASA^3^P directory.

Future directions comprise the development and integration of further analyses, *e.g.* detection and characterization of plasmids, phages and CRISPR cassettes as well as further enhancements in terms of scalability and usability.

## Acknowledgement

The authors thank the BiBiServ team from Bielefeld University for supply of the BiBiGrid framework and especially Jan Krüger for his comprehensive support. Furthermore, we thank Can Imirzalioglu, Jane Falgenhauer, Yancheng Yao and Swapnil Doijad at the Institute of Medical Microbiology of Justus Liebig University Giessen for their ideas, bug reports and fruitful feedback on various aspects regarding implemented analyses and the graphical user interface.

## Supporting information

**S1 Table. Third party executable parameters and options.** Parameters and options without scientific impact are excluded, *e.g.* input/output directories or number of threads.

**S2 Table. Accession numbers of 32 *Listeria monocytogenes* isolates and reference genomes of the ASA^3^P benchmark project.** This exemplary project comprises 32 isolates from SRR Bioproject PRJNA215355 as well as two *Listeria monocytogenes* reference genomes from RefSeq. The project is provided as a GNU zipped tarball at https://s3.computational.bio.uni-giesen.de/swift/v1/asap/example-lmonocytogenes-32.tar.gz

**S3 Table. Information on host CPUs used for wall clock runtime benchmarks.**

**S1 Fig. Exemplary screenshot of configuration template sheet 1.**

**S2 Fig. Exemplary screenshot of configuration template sheet 2.**

**S3 Fig. Exemplary project directory structure.** Each project analyzed by ASA^3^P strictly follows a conventual directory organization and thus forestalls the burden of unnecessary configurations. Shown is an exemplary project structure representing input and output files and directories of the *Listeria monocytogenes* example project. For the sake of readability repeated blocks are collapsed represented by a triple dot ‘…’

**S4 Fig. Wall clock runtimes for varying compute node and isolate numbers.** Runtimes given in hours and separated between comparative and per-isolate internal pipeline stages due to different scalability metrics. Each compute node provides 32 vCPUs and 64 GB memory. *L. monocytogenes* strains were randomly chosen from SRA Bioproject PRJNA215355. (A) Runtimes of a fixed-size compute cluster comprising 4 compute nodes analyzing varying isolate numbers. (B) Runtimes of compute clusters with varying numbers of compute nodes analyzing a fixed amount of 128 isolates.

**S1 File. Comprehensive list of all per-genome key metrics.**

## References

1. Sanger F, Nicklen S, Coulson AR. DNA sequencing with chain-terminating inhibitors. Proc Natl Acad Sci U S A. 1977;74: 5463–5467.

2. van Dijk EL, Auger H, Jaszczyszyn Y, Thermes C. Ten years of next-generation sequencing technology. Trends Genet. 2014;30: 418–426.

3. Goodwin S, McPherson JD, McCombie WR. Coming of age: ten years of next-generation sequencing technologies. Nat Rev Genet. 2016;17: 333–351.

4. Fraser CM, Gocayne JD, White O, Adams MD, Clayton RA, Fleischmann RD, et al. The minimal gene complement of *Mycoplasma genitalium*. Science. 1995;270: 397–403.

5. Fleischmann RD, Adams MD, White O, Clayton RA, Kirkness EF, Kerlavage AR, et al. Whole-genome random sequencing and assembly of *Haemophilus influenzae* Rd. Science. 1995;269: 496–512.

6. Haft DH, DiCuccio M, Badretdin A, Brover V, Chetvernin V, O’Neill K, et al. RefSeq: an update on prokaryotic genome annotation and curation. Nucleic Acids Res. 2018;46: D851–D860.

7. Long SW, Williams D, Valson C, Cantu CC, Cernoch P, Musser JM, et al. A genomic day in the life of a clinical microbiology laboratory. J Clin Microbiol. 2013;51: 1272–1277.

8. Didelot X, Bowden R, Wilson DJ, Peto TEA, Crook DW. Transforming clinical microbiology with bacterial genome sequencing. Nat Rev Genet. 2012;13: 601–612.

9. Tettelin H, Masignani V, Cieslewicz MJ, Donati C, Medini D, Ward NL, et al. Genome analysis of multiple pathogenic isolates of *Streptococcus agalactiae*: Implications for the microbial “pan-genome.” Proceedings of the National Academy of Sciences. 2005;102: 13950–13955.

10. Deurenberg RH, Bathoorn E, Chlebowicz MA, Couto N, Ferdous M, García-Cobos S, et al. Application of next generation sequencing in clinical microbiology and infection prevention. J Biotechnol. Elsevier; 2017;243: 16–24.

11. Review on Antimicrobial Resistance. Tackling Drug-Resistant Infections Globally: final report and recommendations [Internet]. Wellcome Trust; 2016 May. Available: https://amr-review.org/sites/default/files/160525_Final%20paper_with%20cover.pdf

12. Revez J, Espinosa L, Albiger B, Leitmeyer KC, Struelens MJ, ECDC National Microbiology Focal Points and Experts Group ENMFPAE. Survey on the Use of Whole-Genome Sequencing for Infectious Diseases Surveillance: Rapid Expansion of European National Capacities, 2015-2016. Frontiers in public health. Frontiers Media SA; 2017;5: 347.

13. Glaser P, Martins-Simões P, Villain A, Barbier M, Tristan A, Bouchier C, et al. Demography and Intercontinental Spread of the USA300 Community-Acquired Methicillin-Resistant Staphylococcus aureus Lineage. MBio. 2016;7: e02183–15.

14. Holden MTG, Hsu L-Y, Kurt K, Weinert LA, Mather AE, Harris SR, et al. A genomic portrait of the emergence, evolution, and global spread of a methicillin-resistant Staphylococcus aureus pandemic. Genome Res. 2013;23: 653–664.

15. Nübel U. Emergence and Spread of Antimicrobial Resistance: Recent Insights from Bacterial Population Genomics. Curr Top Microbiol Immunol. 398: 35–53.

16. Baur D, Gladstone BP, Burkert F, Carrara E, Foschi F, Döbele S, et al. Effect of antibiotic stewardship on the incidence of infection and colonisation with antibiotic-resistant bacteria and Clostridium difficile infection: a systematic review and meta-analysis. Lancet Infect Dis. 2017;17: 990–1001.

17. Schempp FM, Drummond L, Buchhaupt M, Schrader J. Microbial Cell Factories for the Production of Terpenoid Flavor and Fragrance Compounds. J Agric Food Chem. 2018;66: 2247–2258.

18. Corchero JL, Gasser B, Resina D, Smith W, Parrilli E, Vázquez F, et al. Unconventional microbial systems for the cost-efficient production of high-quality protein therapeutics. Biotechnol Adv. 31: 140–153.

19. Huang C-J, Lin H, Yang X. Industrial production of recombinant therapeutics in *Escherichia coli* and its recent advancements. J Ind Microbiol Biotechnol. 2012;39: 383–399.

20. Baeshen MN, Al-Hejin AM, Bora RS, Ahmed MMM, Ramadan HAI, Saini KS, et al. Production of Biopharmaceuticals in *E. coli*: Current Scenario and Future Perspectives. J Microbiol Biotechnol. 2015;25: 953–962.

21. Wackett LP. Microbial-based motor fuels: science and technology. Microb Biotechnol. 2008;1: 211–225.

22. Zhang W, Yin K, Chen L. Bacteria-mediated bisphenol A degradation. Appl Microbiol Biotechnol. 2013;97: 5681–5689.

23. Singh B, Kaur J, Singh K. Microbial remediation of explosive waste. Crit Rev Microbiol. 2012;38: 152–167.

24. Kip N, van Veen JA. The dual role of microbes in corrosion. ISME J. 2015;9: 542–551.

25. Stephens ZD, Lee SY, Faghri F, Campbell RH, Zhai C, Efron MJ, et al. Big Data: Astronomical or Genomical? PLoS Biol. 2015;13: e1002195.

26. Muir P, Li S, Lou S, Wang D, Spakowicz DJ, Salichos L, et al. The real cost of sequencing: Scaling computation to keep pace with data generation. Genome Biol. Genome Biology; 2016;17: 1–9.

27. Gargis AS, Kalman L, Lubin IM. Assuring the Quality of Next-Generation Sequencing in Clinical Microbiology and Public Health Laboratories. J Clin Microbiol. 2016;54: 2857–2865.

28. Aziz RK, Bartels D, Best AA, DeJongh M, Disz T, Edwards RA, et al. The RAST Server: Rapid Annotations using Subsystems Technology. BMC Genomics. 2008;9: 75.

29. Wattam AR, Davis JJ, Assaf R, Boisvert S, Brettin T, Bun C, et al. Improvements to PATRIC, the all-bacterial Bioinformatics Database and Analysis Resource Center. Nucleic Acids Res. 2017;45: D535–D542.

30. Seemann T. Prokka: Rapid prokaryotic genome annotation. Bioinformatics. 2014;30: 2068–2069.

31. Bolger AM, Lohse M, Usadel B. Trimmomatic: A flexible trimmer for Illumina sequence data. Bioinformatics. 2014;30: 2114–2120.

32. Bankevich A, Nurk S, Antipov D, Gurevich AA, Dvorkin M, Kulikov AS, et al. SPAdes: A New Genome Assembly Algorithm and Its Applications to Single-Cell Sequencing. J Comput Biol. 2012;19: 455–477.

33. Chin C-S, Alexander DH, Marks P, Klammer AA, Drake J, Heiner C, et al. Nonhybrid, finished microbial genome assemblies from long-read SMRT sequencing data. Nat Methods. Nature Publishing Group, a division of Macmillan Publishers Limited. All Rights Reserved.; 2013;10: 563–569.

34. Wick RR, Judd LM, Gorrie CL, Holt KE. Unicycler: Resolving bacterial genome assemblies from short and long sequencing reads. Phillippy AM, editor. PLoS Comput Biol. Public Library of Science; 2017;13: e1005595.

35. Bosi E, Donati B, Galardini M, Brunetti S, Sagot MF, Lió P, et al. MeDuSa: A multi-draft based scaffolder. Bioinformatics. 2015;31: 2443–2451.

36. Jia B, Raphenya AR, Alcock B, Waglechner N, Guo P, Tsang KK, et al. CARD 2017: Expansion and model-centric curation of the comprehensive antibiotic resistance database. Nucleic Acids Res. 2017;45: D566–D573.

37. Chen L, Zheng D, Liu B, Yang J, Jin Q. VFDB 2016: Hierarchical and refined dataset for big data analysis - 10 years on. Nucleic Acids Res. 2016;44: D694–D697.

38. Goris J, Konstantinidis KT, Klappenbach JA, Coenye T, Vandamme P, Tiedje JM. DNA-DNA hybridization values and their relationship to whole-genome sequence similarities. Int J Syst Evol Microbiol. 2007;57: 81–91.

39. Wood DE, Salzberg SL. Kraken: ultrafast metagenomic sequence classification using exact alignments. Genome Biol. 2014;15: R46.

40. Camacho C, Coulouris G, Avagyan V, Ma N, Papadopoulos J, Bealer K, et al. BLAST+: architecture and applications. BMC Bioinformatics. 2009;10: 421.

41. Cole JR, Wang Q, Fish JA, Chai B, McGarrell DM, Sun Y, et al. Ribosomal Database Project: data and tools for high throughput rRNA analysis. Nucleic Acids Res. 2014;42: D633–42.

42. Kurtz S, Phillippy A, Delcher AL, Smoot M, Shumway M, Antonescu C, et al. Versatile and open software for comparing large genomes. Genome Biol. 2004;5: R12.

43. Jolley KA, Bray JE, Maiden MCJ. A RESTful application programming interface for the PubMLST molecular typing and genome databases. Database. 2017;2017. doi:10.1093/database/bax060

44. Langmead B, Salzberg SL. Fast gapped-read alignment with Bowtie 2. Nat Methods. 2012;9: 357–359.

45. Li H. Minimap2: pairwise alignment for nucleotide sequences. Bioinformatics. 2018;34: 3094–3100.

46. Li H, Handsaker B, Wysoker A, Fennell T, Ruan J, Homer N, et al. The Sequence Alignment/Map format and SAMtools. Bioinformatics. 2009;25: 2078–2079.

47. Cingolani P, Platts A, Wang LL, Coon M, Nguyen T, Wang L, et al. A program for annotating and predicting the effects of single nucleotide polymorphisms, SnpEff. Fly. 2012;6: 80–92.

48. Price MN, Dehal PS, Arkin AP. FastTree 2--approximately maximum-likelihood trees for large alignments. PLoS One. 2010;5: e9490.

49. Page AJ, Cummins CA, Hunt M, Wong VK, Reuter S, Holden MTG, et al. Roary: Rapid large-scale prokaryote pan genome analysis. Bioinformatics. 2015;31: 3691–3693.

